# The industrial solvent trichloroethylene induces LRRK2 kinase activity and dopaminergic neurodegeneration in a rat model of Parkinson’s disease

**DOI:** 10.1101/2020.11.02.365775

**Authors:** Sandra L. Castro, Emily M. Rocha, Christopher R. Bodle, Katrina E. Johnson, J. Timothy Greenamyre, Briana R. De Miranda

**Author notes:** Correspondence should be addressed to Briana R. De Miranda, University of Alabama at Birmingham, 1719 6^th^ Ave South | CIRC 560 | Birmingham, AL 35924.

## Abstract

Gene-environment interaction is implicated in the majority of idiopathic Parkinson’s disease (PD) risk, and some of the most widespread environmental contaminants are selectively toxic to dopaminergic neurons. Pesticides have long been connected to PD incidence, however, it has become increasingly apparent that other industrial byproducts likely influence neurodegeneration. For example, organic solvents, which are used in chemical, machining, and dry-cleaning industries, are of growing concern, as decades of solvent use and their effluence into the environment has contaminated much of the world’s groundwater and soil. Like some pesticides, certain organic solvents, such as the chlorinated halocarbon trichloroethylene (TCE), are mitochondrial toxicants, which are collectively implicated in the pathogenesis of dopaminergic neurodegeneration. Recently, we hypothesized a possible gene-environment interaction may occur between environmental mitochondrial toxicants and the protein kinase LRRK2, mutations of which are the most common genetic cause of familial and sporadic PD. In addition, emerging data suggests that elevated wildtype LRRK2 kinase activity also contributes to the pathogenesis of idiopathic PD. To this end, we investigated whether chronic, systemic TCE exposure (200 mg/kg) in aged rats produced wildtype LRRK2 activation and influenced predegenerative dopaminergic dysfunction. Interestingly, we found that TCE not only induced LRRK2 kinase activity in the brain, but produced a significant dopaminergic lesion in the nigrostriatal tract, elevated oxidative stress, and caused endolysosomal dysfunction and protein accumulation (α-synuclein). Together, these data suggest that TCE-induced LRRK2 kinase activity contributed to the selective toxicity of dopaminergic neurons. We conclude that gene-environment interactions between certain industrial contaminants and LRRK2 likely influence PD risk.

## Introduction

Halogenated organic solvents are heavily used chemicals throughout the world and represent widespread industrial byproducts that contaminate the environment^1–4^. The chlorinated solvent trichloroethylene (TCE) is a common degreasing product and chemical feedstock, which was widely used during the 20^th^ century and remains in regulated use today^5,6^. Due to its chlorinated structure, TCE does not readily break down in the environment, resulting in lasting contamination of the solvent, particularly in ground water^7^. In recent decades TCE has been detected in up to 34% of drinking water, and is currently estimated to contaminate the treated water supply of 14-20 million individuals within the United States^8^. Despite its continued use, TCE is a well-known human toxicant, implicated in renal cancer, liver cancer, non-Hodgkin’s lymphoma, autoimmune disease, and neurotoxicity^9^. TCE is also implicated in the development of the most common neurodegenerative movement disorder, Parkinson’s disease (PD), as some evidence suggests occupational exposure to TCE is associated with increased PD risk^10,11^. Similarly, proof-of-principal neurotoxicity studies describe the degeneration of dopaminergic neurons following high levels (800-1000 mg/kg) of TCE treatment in rodents^12,13^. Thus, TCE represents a candidate environmental or occupational toxicant that could increase risk for PD, however, it is unclear what mechanisms are responsible for its selective neurotoxicity to dopaminergic neurons.

The vast majority – about 85% – of PD is considered idiopathic (iPD), and represents a spectrum of disease sequelae among patients, including motor and non-motor symptoms that progressively worsen^14^. For most, gene-environment interaction (GxE) likely drives the majority of iPD risk, however, quantifiable GxE is difficult to assess^15^. The most commonly inherited genetic mutations associated with elevated risk for PD are within the leucine rich repeat kinase 2 (LRRK2) gene, and account for approximately 1-2% of PD cases^16^. Recently, we and others have observed that wildtype LRRK2 may play a role in iPD^17,18^, as elevated LRRK2 protein kinase activity was observed in dopaminergic neurons of postmortem brain tissue^17^, peripheral blood cells^19^, and urinary exosomes of iPD patients ^20^. While the cause of aberrant LRRK2 kinase activity in iPD is unknown, endogenous, wildtype LRRK2 appears to be induced, either directly or indirectly by oxidative stress^21^. A potential source for this is the cellular oxidative damage produced by mitochondrial toxicants (e.g. pesticides), which are implicated as environmental factors in PD risk^22–24^.

LRRK2 is a master regulator of protein and vesicular trafficking; its kinase substrates, Rab GTPases, drive the direction and fusion of endosomal vesicles important to the autophagy-lysosomal pathway (ALP)^25^. Increases in LRRK2 kinase activity, whether the result of inherited mutations (e.g. G2019S) or overactivation of the wildtype protein, results in ALP dysfunction^26^, and is implicated in the accumulation of protein (e.g. α-synuclein)^27–29^. As such, endolysosomal impairment mediated by LRRK2 appears to be an early and important pathological factor in dopaminergic cell survival. As a mitochondrial toxicant^30^, TCE represents a potential environmental source for LRRK2 kinase activation that could induce a cascade of predegenerative pathology within dopaminergic neurons, ultimately resulting in neurodegeneration over time.

To this end, we have investigated whether a gene-environment interaction exists between the ubiquitous environmental contaminant TCE and LRRK2, which could provide a mechanism for the observed dopaminergic pathology following TCE exposure. To identify TCE-induced LRRK2 kinase activity and associated neuropathology, we developed a rat model of moderate TCE exposure (200 mg/kg) supplied in a daily gavage over a chronic dosing period (6 weeks) in aged Lewis rats. As this dose is considerably lower than previously reported studies, the basis of pathological analysis for this study was to identify predegenerative changes within nigrostriatal dopaminergic neurons, potentially as a result of TCE-induced LRRK2 kinase activity. Surprisingly, we observed a marked loss of dopaminergic neurons and their striatal terminal projections, corresponding with a reduction in motor behavior following 6-week, 200 mg/kg TCE exposure in aged rats. Within surviving dopaminergic neurons, TCE caused a clear increase in oxidative stress, produced dysfunction in endolysosomal and protein degradation pathways, and resulted in the accumulation of endogenous α-synuclein. TCE also caused LRRK2 kinase activation in the nigrostriatal tract, detected by protein auto- and substrate phosphorylation, which occurred prior to any significant dopaminergic neurodegeneration. Collectively, these data show that a moderate, *systemic* TCE exposure resulted in elevated LRRK2 kinase activity in the brain that mirrors pathological activation observed in PD patients, providing an avenue for PD risk from exposure to this industrial contaminant.

## Materials and Methods

### Chemical reagents and supplies

Trichloroethylene (CAS Number 79-01-6) and other chemicals were purchased from Sigma-Aldrich (St. Louis, MO) unless otherwise noted. Antibody information for immunohistochemistry (IHC), immunocytochemistry (ICC) and western blot is listed in **Table 1**.

**Table 1.** Primary antibodies.

| Antigen | Antibody Catalog Information | Company | IHC or Western Blot Concentration |
| --- | --- | --- | --- |
| Tyrosine hydroxylase | AB1542 | EMD Millipore (Burlington, MA) | 1:2000 |
| **IHC**<br>Ser129-α-Synuclein<br>Lamp1 | ab51253<br>ab24170 | Abcam (Cambridge, MA) | 1:500<br>1:500 |
| **Western Blot**<br>pThr73-Rab10<br>pSer935-LRRK2<br>pSer1292-LRRK2<br>LRRK2 c41-2 | ab231707<br>ab133450<br>ab203181<br>ab133474 |  | 1:2000<br>1:2000<br>1:2000<br>1:2000 |
| **ICC**<br>pThr73-Rab10 | ab241060 |  | 1:1000 |
| Iba1 | 019-19741 | Wako Chemicals USA (Irvine, CA) | 1:500 |
| CD68 (ED1) | MCA341 | BioRad (Hercules, CA) | 1:500 |
| α-Synuclein | 610787 | BD Biosciences (San Jose, CA) | 1:500 |
| p62/SQSTM1 (2C11) | H00008878-M01 | Abnova (Tapei, Taiwan) | 1:500 |
| **IHC**<br>pThr73-Rab10 | 10<sup>th</sup> bleed | MRC Protein Unit (Dr. Dario Alessi, University of Dundee) | 1:500 |
| **IHC**<br>Total LRRK2 | N241A/34 | NeuroMab UC Davis (Davis, CA) | 1:500 |
| **IHC**<br>14-3-3 | NBP2-67831 | Novus Biologics (Centennial, CO) | 1:500 |

### Animals and TCE administration

Three separate cohorts of rats were administered TCE in this study (**Supplemental Figure 1**). Adult, male and female Lewis rats were obtained through a retired breeding program from Envigo (Indianapolis, IN). Upon arrival, rats were separated into single-housing two weeks prior to the onset of treatment and handled daily to prevent stress upon study commencement. Conventional diet and water were available to rats *ad libitum,* and animals were maintained under standard temperature-controlled conditions with a 12-hour light-dark cycle. TCE was dissolved into premium olive oil (Trader Joe’s, Monrovia, CA) for a final concentration of 200 mg/kg based on individual rat weight (cohorts 1 and 2), or 50, 100, 200, 400 or 800 mg/kg (Cohort 3). TCE and vehicle (olive oil) groups were randomly divided, and each animal was administered a single daily oral gavage for one, three, or six weeks corresponding to each Cohort endpoint. Animal weight was recorded daily, and rats were observed twice a day for signs of overt toxicity or animal morbidity (**Supplemental Figure 1**). Following TCE exposure, animals were euthanized using a lethal dose of pentobarbital, followed by transcardial perfusion with PBS and fixation with 4% paraformaldehyde (Cohorts 1 and 2). Cohort 3 rat brain tissue was collected, flash frozen, and stored at −80°C. Following euthanasia and tissue collection, animals were assigned a four-digit code and researchers were blind to treatment groups during all subsequent analyses. All experiments involving animal treatment and euthanasia were approved by the University of Pittsburgh Institutional Animal Care and Use Committee. TCE handling and disposal was carried out following University of Pittsburgh Environmental Health and Safety procedures.

### Motor behavior

Open field behavior was monitored after 6 weeks of TCE treatment (200 mg/kg) in Cohort 1, aged male Lewis rats. Following a period of acclimation, each rat was placed in an open field chamber and allowed to freely explore for 20 minutes while tracking software (Noldus EthoVision XT, Leesburg, VA) recorded a standard array of open field parameters. Motor behavior analyses were carried out following all protocols set by the University of Pittsburgh Rodent Behavioral Analysis Core (RBAC), including ambient noise, light, and animal handling parameters to limit rodent stress and erroneous stimuli that could affect behavioral outcomes.

### Striatal terminal intensity

A series of brain sections (35 μm) encompassing the volume of the rat striatum (1/6 sampling fraction, approximately 10 sections per animal) were stained for tyrosine hydroxylase (TH) and detected using an infrared secondary antibody (IRDye® 680, LiCor Biosciences). Striatal tissue sections were analyzed using near-infrared imaging for density of dopamine neuron terminals (LiCor Odyssey), and quantified using LiCor Odyssey software (V3.0; Licor Biosciences, Lincoln, NE) with consistent background subtraction. Results are reported as striatal TH intensity in arbitrary fluorescence units.

### Stereology

Stereological analysis of dopamine neuron number in the SN was achieved using an adapted protocol from Tapias et al.^31,32^ as reported in De Miranda et al.^33,34^ employing an unbiased, automated system. Briefly, nigral tissue sections were stained for TH and counterstained with DAPI and Nissl NeuroTrace Dye (640; Life Technologies) then imaged using a Nikon 90i upright fluorescence microscope equipped with high N.A. plan fluor/apochromat objectives, Renishaw linear encoded microscope stage (Prior Electronics) and Q-imaging Retiga cooled CCD camera (Center for Biological Imaging, University of Pittsburgh). Images were processed using Nikon NIS-Elements Advanced Research software (Version 4.5, Nikon, Melville, NY), and quantitative analysis was performed on fluorescent images colocalizing DAPI, TH, and Nissl-positive stains. Results are reported as the number of TH-positive cell bodies (whole neurons) within the SN.

### Immunohistochemistry and immunopathology

Brain sections (35 μm) were maintained at −20°C in cryoprotectant, stained while free-floating, and mounted to glass slides for imaging, using a “primary antibody delete” (secondary antibody only) stained section to subtract background fluorescence. The proximity ligation assay (PLA) was performed using the DuoLink (Millipore Sigma, St. Louis, MO) protocol, and as reported in Di Maio et al. (2018)^17^. Briefly, fixed rat brain tissue was stained for TH (Ab1542, Millipore) followed by staining with total LRRK2 (N241A/34, NeuroMab, UC Davis), and 14-3-3 (ab9063, Abcam) mouse and rabbit primary antibodies, respectively, overnight at 4°C. DuoLink rabbit plus and mouse minus probes conjugated to oligonucleotides (Step 1) were incubated for 1 hour at 37°C in a humid chamber. Ligation of the probes using DuoLink Ligase Solution (Step 2) occurred for 30 min at 37°C in a humid chamber. Amplification of the probes using DuoLink Green (488) Detection Reagents (Step 3) was achieved by a 100 minute incubation at 37°C in a humid chamber. Following PLA amplification, tissue sections were washed and coverslipped and imaged 24-hours later.

Fluorescent immunohistochemical images were collected using an Olympus BX61 confocal microscope and Fluoview 1000 software (Melville, NY). Quantitative fluorescence measurements were thoroughly monitored using standard operating imaging parameters to ensure that images contained no saturated pixels. For quantitative comparisons, all imaging parameters (e.g., laser power, exposure, pinhole) were held constant across specimens. Confocal images were analyzed using Nikon NIS-Elements Advanced Research software (Version 4.5, Nikon, Melville, NY). A minimum of 6 images per tissue slice were analyzed per animal, averaging 9-15 neurons per 60-100x image (approximately 120-180 cells per animal, per histological stain depending on magnification). 20x magnification was used to generate montage imaging of the ventral midbrain or striatum, for representative images or analysis per image using anatomical region of interest (ROI) boundaries. Results are reported as a measure of puncta within TH-positive cells, either number of objects (# of objects), area (μm^2^), or fluorescence intensity (arbitrary units).

### Western blots

Brain tissue from Cohort 3 rats (3-week TCE treatment and dose response) was microdissected and flash frozen. Tissue lysates were prepared by homogenizing each brain section in a Dounce tissue grinder using 20% weight/volume lysis buffer (see below for recipe). Tissue was ground using 20 loose strokes and 20 tight strokes, then left on ice for 30 minutes to lyse. Homogenized tissue was centrifuged at 15,000xg for 10 minutes at 4°C, and supernatant was collected and centrifuged for an additional 15,000xg for 10 minutes at 4°C to pellet remaining debris. Protein was quantified using a Pierce Bovine Serum Albumin Standard (23208, ThermoFisher Scientific). 10x Cell Signaling Lysis Buffer (9803, Danvers, MA) was diluted to 1x using ultra pure water, followed by the addition of a protease inhibitor cocktail (#78430) and phosphatase inhibitor cocktail (788427) from ThermoFisher Scientific (Waltham, MA).

Samples were incubated with NuPage LDS 4x sample buffer (NP0007, ThermoFisher Scientific) and boiled for 10 minutes at 95°C. A total of 100 μg brain lysate was run on a Bio-Rad Mini-Protean TGX 4-20% polyacrylamide gel (4561094, Bio-Rad, Hercules, CA) at 200V for approximately 40 minutes. Gels were transferred to a nitrocellulose membrane (1704159, Bio-Rad) using the Bio-Rad Trans-Blot Turbo system for 7 minutes. Nitrocellulose membranes were blocked for 1 hour in 50% TBS (20mM Tris, 0.5M NaCl, pH 7.5) 50% Odussey blocking buffer (LiCor, Lincoln, NE), then incubated overnight with primary antibodies in 50% TBS-T (20 mM Tris, 0.5 M NaCl, pH 7.5, 0.1% Tween 20), 50% Odyssey blocking buffer at 4°C. Membranes were washed 3 x 10 minutes in TBS-T followed by secondary incubation with Odyssey IRDye® 800 (926-32213) and IRDye® 680 (926-68072) for 1 hour at room temperature. Following 3 x 10 minute wash in TBS-T, membranes were scanned using the Odyssey CLx imaging system, and analyzed using LiCor Image Studio Software. Quantified intensity is reported as phosphorylated protein in relation to total LRRK2, in relation to β-actin.

### Blood cell analysis

Rodent blood was collected via cardiac stick in Cohorts 1-3. Approximately 1000 μl blood was extracted and transferred to K2 EDTA tubes (Thermo Fisher, Waltham MA). Samples were spun at 2500g for 20 minutes and the plasma was removed and stored. The buffy coat enriched fraction was collected and diluted in 1000 μl OptiMEM reduced serum medium (Thermo Fisher). Samples were then plated at 100 μl per well in 16 well chamber slides, which were pre-coated with Cell-Tak (Corning Inc, Corning NY). Chamber slides were centrifuged at 500g for 5 min to promote cell adherence. Post centrifuge, media was removed, and cells were fixed with 4% PFA for 20 min at RT. Samples containing white blood cells (WBC) were stained using pThr73-Rab10 (ab241060, Abcam) overnight at 4°C, and analyzed via confocal microscopy as described above.

### Statistical analyses

All data are expressed as mean values ± standard error of the mean (SEM). For statistical comparison between vehicle and TCE groups, a non-parametric T-test was used. Statistical significance between male and female rats was evaluated for normally distributed means by two-way analysis of variance (two-way ANOVA) with a Sidak multiple comparisons test to correct for mean comparison between multiple groups, where source of variation was defined as “sex” or “TCE” groups. P-value and F-statistic are reported for the interaction comparison between “vehicle” and “TCE” variables as F (DFn, DFd) = x within each figure legend. Prior to study onset, *a-priori* power analyses were conducted using G*power software (Heinrich-Heine-University Düsseldorf) to determine the sample size required for a 20-40% difference between mean, with a 95% power at α = 0.05. Statistical significance between treatment groups is represented in each figure as \**p* < 0.05, \*\**p*<0.01, \*\*\**p*<0.001, \*\*\*\**p*<0.0001, unless otherwise specified on graph. Statistical outliers from each data set were determined using the extreme studentized deviate (Grubbs’ test, *α* = 0.05). Statistical analyses were carried out using GraphPad Prism software (V. 5.01).

## Results

### Aged rats exposed to TCE display selective nigrostriatal neurodegeneration

Aged adult (15 month), male Lewis rats were given a daily oral gavage of 200 mg/kg TCE or vehicle (olive oil) for 6-weeks (Cohort 1, Figure 1a). Following 6-weeks of TCE exposure, locomotor behavior was assayed using the Open Field test, which revealed a significant reduction in total Distance Traveled, as well as Ambulatory Time in rats receiving TCE treatment (Figure 1b-d). The reduction in spontaneous locomotor activity was also evident from trace plots of TCE-treated animals from open field monitoring (Figure 1e). Immunohistochemistry of striatal brain sections from TCE-treated rats revealed approximately a 30% reduction in tyrosine hydroxylase (TH) intensity within the striatum compared to vehicle (*p* = 0.0001), indicative of a loss of dopaminergic neuron terminals (Figure 1f-g). Assessment of neuronal subtypes within the striatum, such as DARPP32-positive medium spiny neurons and GAD67-positive neurons, showed no remarkable change in protein expression. Similarly, total neuronal cell bodies within the striatum visualized using NeuN did not appear altered between vehicle and TCE treated rats (Supplemental Figure 2a).

**Figure 1.**
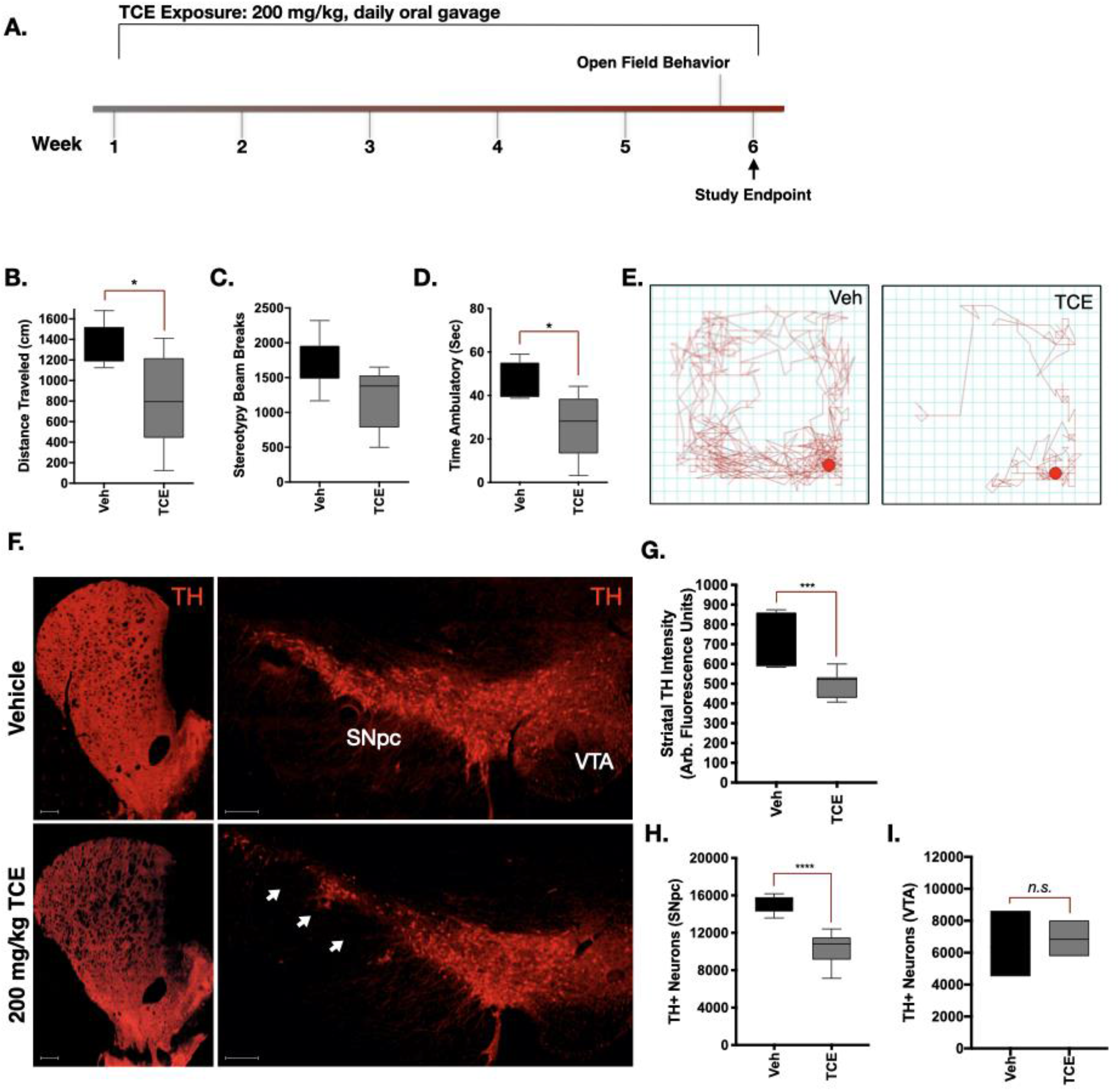
Aged rats exposed to TCE display selective nigrostriatal neurodegeneration. **A.** 15-Month-old, male Lewis rats were exposed to 200 mg/kg TCE or vehicle (olive oil) via daily oral gavage for 6 weeks. **B-D.** Quantitative parameters measured from open-field chamber motor behavior analysis showed significant differences in “Distance Traveled” (F(4, 5 = 5.267), *p* = 0.015) and “Time Ambulatory” (F(4, 4 = 3.182), *p* = 0.03) but not “Beam Breaks” (F(4, 5 = 1.408), *p* = 0.06). **E.** Depiction of a trace plot from free movement of TCE or vehicle treated rats after 6 weeks. **F.** Representative 20x montage images of tyrosine hydroxylase (TH; red) positive cells in the striatum, substantia nigra pars compacta (SNpc), and ventral tegmental area (VTA). **G-I.** Quantification of dopaminergic cell loss from the striatum (F(5, 11 = 3.567), *p* = 0.0001) and SNpc (F(8, 4 = 2.959), *p*<0.0001) but not the VTA (F(2, 2 = 3.46) (*p* = 0.73) following 6 weeks of TCE exposure. Statistical analysis unpaired T-test, error bars represent SEM, (N = 7 vehicle, 10 TCE).

TCE caused approximately 32% loss of dopaminergic neurons in the substantia nigra (SN) assessed by stereology, compared to vehicle (*p*<0.0001; Figure 1f,h). Dopaminergic neurons within the ventral tegmental area (VTA) were not significantly altered by TCE exposure in these animals (*p* = 0.73; Figure 1f,i). Stereological TH counts were colocalized with Nissl stain for dopaminergic cell bodies within the SN, which show the visual loss of cell volume following TCE treatment (Supplemental Figure 2b).

### TCE induced oxidative stress within dopaminergic neurons

Evidence of oxidative stress was measured using markers for oxidative protein modification within the dopaminergic neurons of the SN from TCE or vehicle treated rats. Significantly elevated levels of 3-nitrotyrosine (3-NT; *p*<0.001), a marker for peroxynitrite (ONOO^−^) protein adducts on tyrosine residues were observed in dopaminergic (TH-positive) neurons of TCE-exposed aged rats (Figure 2a,b). Similarly, elevated levels of the protein 4-hydroxynonenal (4-HNE; *p*<0.01), a marker for lipid peroxidation caused by oxygen radicals, were observed in dopaminergic neurons of the SN within TCE treated rats (Figure 2a,c). Together, these data indicate surviving dopaminergic neurons within TCE treated animals display evidence of elevated levels of oxidative damage compared to age-matched, vehicle treated rats.

**Figure 2.**
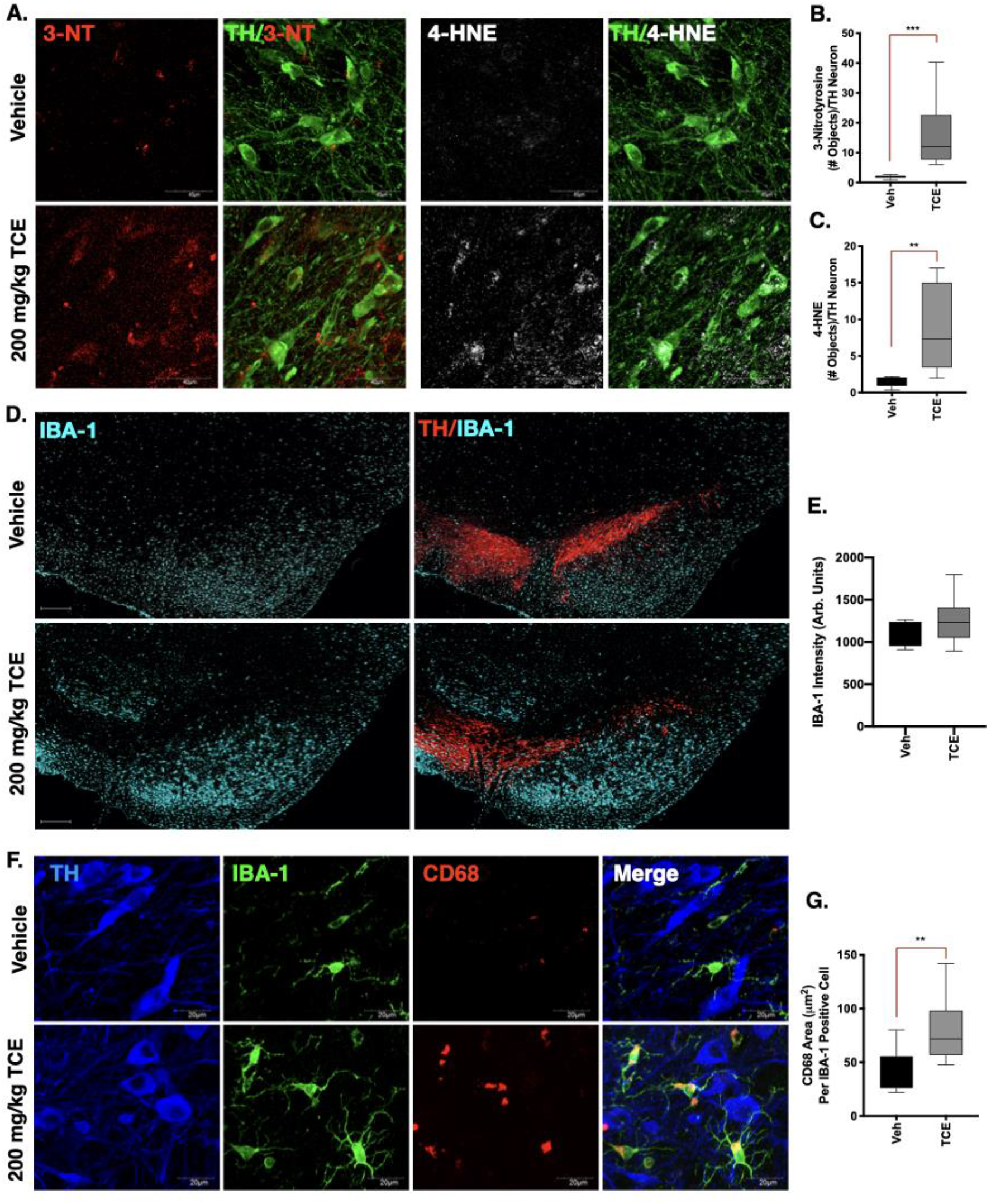
Evidence of oxidative damage and neuroinflammation in the ventral midbrain of TCE exposed rats. Immunohistochemistry was conducted in brain tissue from 15-month old male Lewis rats exposed to 200 mg/kg TCE or vehicle (olive oil) for 6 weeks. **A.** Representative confocal images (60x) of the peroxynitrite adduct 3-nitrotyrosine (3-NT, red) and the lipid peroxidation maker 4-hydroxynonenal (4-HNE, white) in dopaminergic neurons of the SNpc (TH, green). **B-C.** Quantification of 3-NT (F(8, 10 = 517.6), *p* = 0.0006) and 4-HNE (F(8, 6 = 65.19), *p* = 0.0061) within dopaminergic neurons indicates elevated oxidative damage following 6 weeks of TCE exposure. **D.** Representative images (20x montage) of microglia (IBA-1, cyan) and dopaminergic neurons (TH, red) in the SNpc of vehicle and TCE treated rats. **E.** Quantification of IBA-1 intensity is not significantly different (F(13, 3 = 2.82), *p* = 0.309) between treatment groups. **F.** Representative confocal images (60x) of IBA-1 (green) and the myeloid lysosomal protein CD68 (red), surrounding dopaminergic neurons (TH, blue) in the SNpc of TCE or vehicle treated rats. **G.** Significantly elevated CD68 area in the SNpc of TCE treated rats compared to vehicle (F(13, 6 = 1.59), *p* = 0.005). Statistical analysis unpaired T-test, error bars represent SEM, (N = 7 vehicle, 10 TCE).

### Moderate TCE exposure results in a mildly activated microglial phenotype

Previous studies using high doses of TCE exposure in rats (up to 1,000 mg/kg) reported a hypertrophic microglial phenotype consistent with activated microglia ^13^. Despite the lower dose used here (200 mg/kg), we observed a similar trend in hypertrophic microglia throughout the ventral midbrain and surrounding the SN in TCE treated rats compared to vehicle (Figure 2d). Observational differences in IBA-1 positive cell volume, however, were not statistically different between treatment groups (*p* = 0.309; Figure 2c). Conversely, high-resolution analysis of lysosomal protein CD68 area within IBA-1 positive cells, indicative of increased ongoing phagocytic activity, was significantly elevated (*p*<0.001) in the microglia of TCE treated rats (Figure 2e-f).

### Endolysosomal dysfunction and protein accumulation in the SN of TCE-treated rats

Within nigral dopaminergic neurons, we observed a marked increase of Rab5 positive puncta in rats exposed to TCE, an indication that early endosomes were accumulating within these cells (*p*<0.01; Figure 3a,b). TCE exposure also caused a significant accumulation of protein p62/Sequestosome 1 within dopaminergic neurons of the SN, the ubiquitin-like protein that tags cargo destined for lysosomal degradation (*p*<0.05; Figure 3a,c). In conjunction, a robust decrease of the late endosome and lysosomal membrane protein Lamp1 was observed in dopaminergic neurons of the SN in TCE treated rats (p<0.0001; Figure 3a,d). Collectively, these data suggest TCE exposure caused impairment of ALP within the surviving nigral dopaminergic neuron population. These changes also correlated with accumulation of both phosphorylated (pSer129-αSyn; *p* = 0.0076) and total α-synuclein (*p* = 0.0003) within the soma of dopaminergic neurons of TCE treated animals (Figure 3e-g).

**Figure 3.**
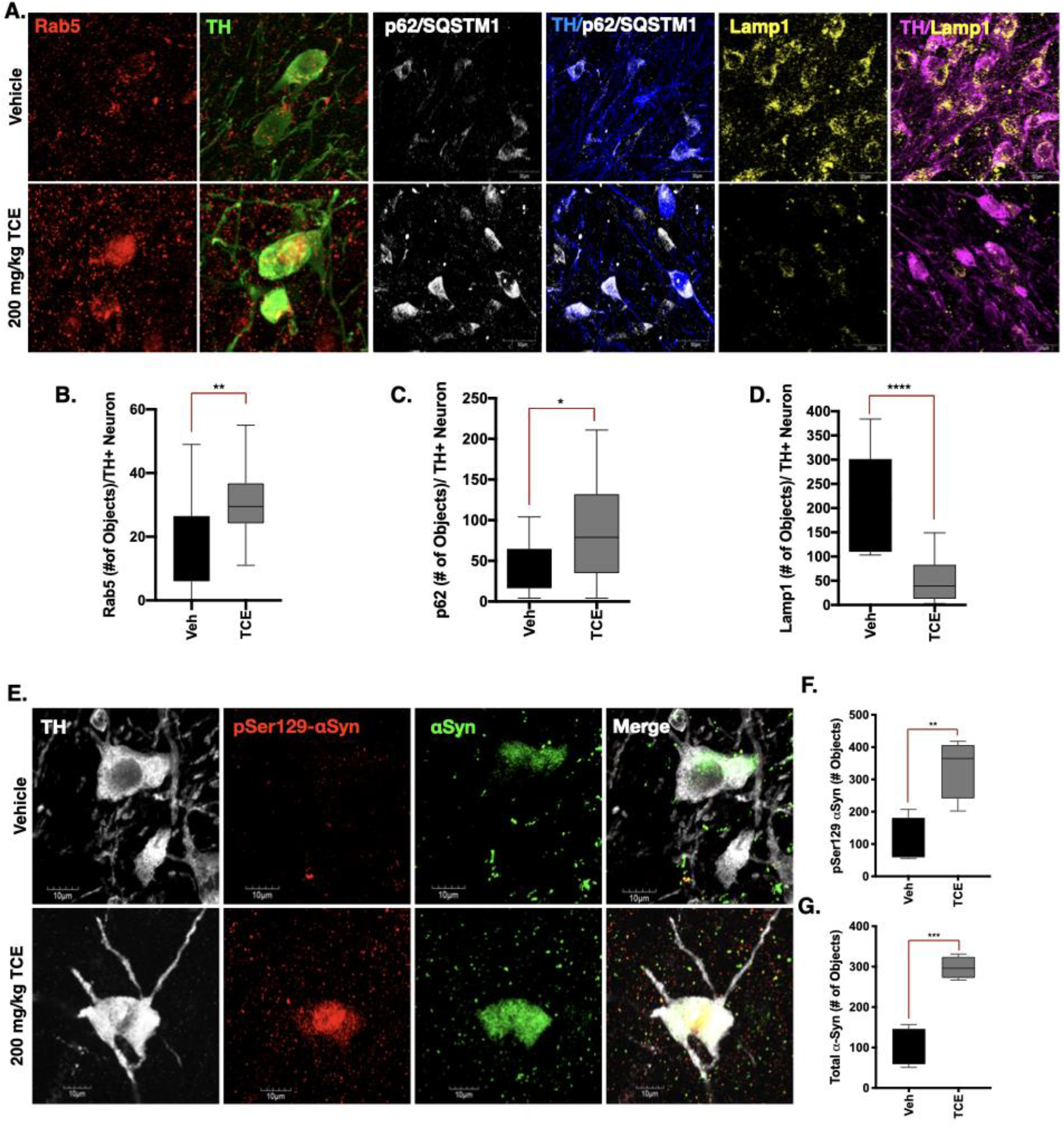
Endolysosomal dysfunction and protein accumulation occurs in dopaminergic neurons of TCE-treated rats. Immunohistochemistry was conducted in brain tissue from 15-month old male Lewis rats exposed to 200 mg/kg TCE or vehicle (olive oil) for 6 weeks. **A.** Representative confocal images of Rab5 (200x, red), p62/SQSTM1 (100x, white), and Lamp1 (100x, yellow) in the SNpc of vehicle or TCE treated rats (counterstained for TH, green, blue, magenta, respectively). **B.** The early endosome marker Rab5 is significantly elevated in dopaminergic neurons in the SNpc of rats exposed to TCE (F(12, 23 = 1.62), *p* = 0.0087). **C.** The ubiquitin like protein p62/SQSTM1 is also significantly elevated in TCE treated rats (F(14, 9 = 3.976), *p* = 0.035). **D.** The lysosomal membrane protein Lamp1 is significantly reduced in dopaminergic neurons of rats treated with TCE (F(9, 14 = 4.84), *p*<0.0001). **E-G.** Phosphorylated (pSer129-αSyn, red) and total αSyn (green) protein was accumulated in dopaminergic neurons (TH, white) of rats exposed to TCE; pSer129-αSyn (F(3, 3 = 1.87), *p* = 0.0076), total αSyn (F(3, 3 = 2.82), *p* = 0.0003). Statistical analysis unpaired T-test, error bars represent SEM, (N = 7 vehicle, 10 TCE).

### TCE exposure influences LRRK2 protein-protein interactions associated with elevated kinase activity

A proximity ligation (PL) assay was performed in rat brain tissue between LRRK2 and its regulatory protein 14-3-3, which is inversely correlated with LRRK2 kinase activity when 14-3-3 is bound at phosphorylation sites Ser910 and Ser953 (N-terminal) or Ser1444 (Roc domain^35–38^ Within dopaminergic neurons, PL signal between LRRK2 and 14-3-3 was significantly lower (*p*<0.008) in TCE treated rats than vehicle, providing evidence that LRRK2 kinase activity was elevated in these cells (Figure 4a-b). In conjunction, Rab10, a kinase substrate of LRRK2, was highly phosphorylated (pThr73) in dopaminergic neurons (*p*<0.05) in the SN of TCE treated rats Figure 4c-d, f-g). Six weeks of TCE exposure also resulted in elevated pThr73-Rab10 signal compared to vehicle in white blood cells (WBC) cultured from rat blood samples collected at study termination (*p*<0.05; Figure 4e).

**Figure 4.**
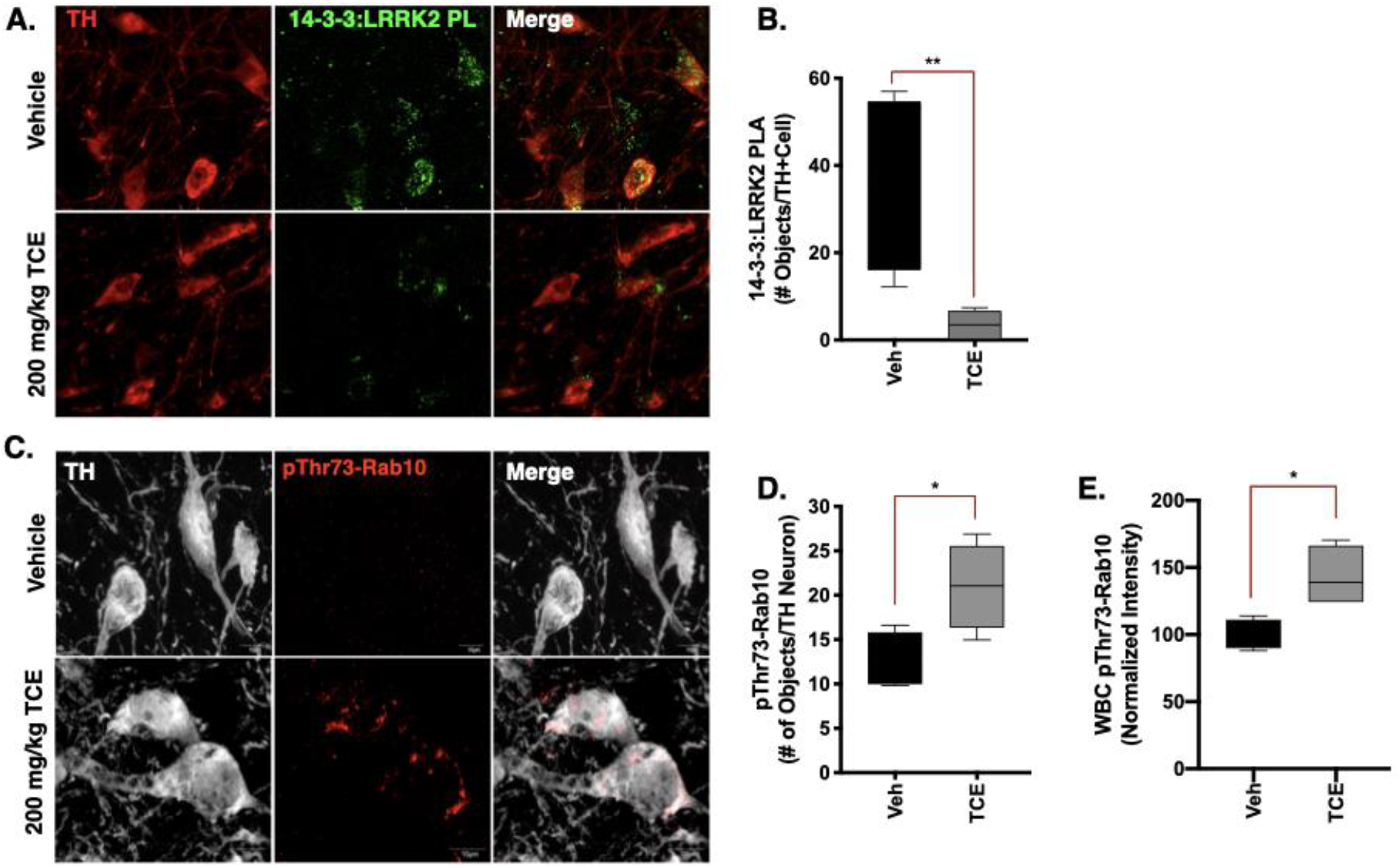
TCE exposure influences LRRK2 protein-protein interactions associated with elevated kinase activity. Immunohistochemistry was conducted in brain tissue from 15-month old male Lewis rats exposed to 200 mg/kg TCE or vehicle (olive oil) for 6 weeks. **A-B.** Proximity ligation assay (PLA) between LRRK2 (N241A) and its regulatory protein 14-3-3 (PL; green) in dopaminergic neurons (TH, red) of TCE treated animals shows significant reduction in signal compared to vehicle (F(3, 4 = 35.43), *p* = 0.0084). **C-E.** An antibody generated towards pThr73-Rab10 (red) indicates a significant accumulation of the LRRK2 kinase substrate in dopaminergic neurons (TH, white; F(3, 3 = 1.87), *p* = 0.0076), and in white blood cells (WBC) cultured from Lewis rats after 6 weeks of TCE exposure (F(3, 3 = 4.20), *p* = 0.0238). Statistical analysis unpaired T-test, error bars represent SEM, (N = 7 vehicle, 10 TCE).

### TCE-induced LRRK2 kinase activity is elevated prior to dopaminergic neuron loss

To assess the temporal loss of dopaminergic neurons in response to TCE exposure, adult male and female rats were treated for one, three, or six weeks with 200 mg/kg TCE (or vehicle, olive oil) via daily oral gavage (Cohort 2, Figure 5a). Analysis of striatal TH intensity over the time course, revealed progressive dopaminergic terminal loss from three-weeks to six-weeks of continuous TCE exposure in both male (*p*<0.001) and female animals (*p*<0.0001; Figure 5b). Striatal terminal loss induced by TCE preceded dopaminergic neuron death, as no significant neurodegeneration was observed until six weeks of TCE exposure (*p* = 0.005; Figure 5c). In contrast to other neurotoxicant models, such as rotenone ^34^, it did not appear that sex influenced a robust difference in dopaminergic neurodegeneration in TCE treated rats (*p* = −6795, two-way ANOVA, α = 0.05).

**Figure 5.**
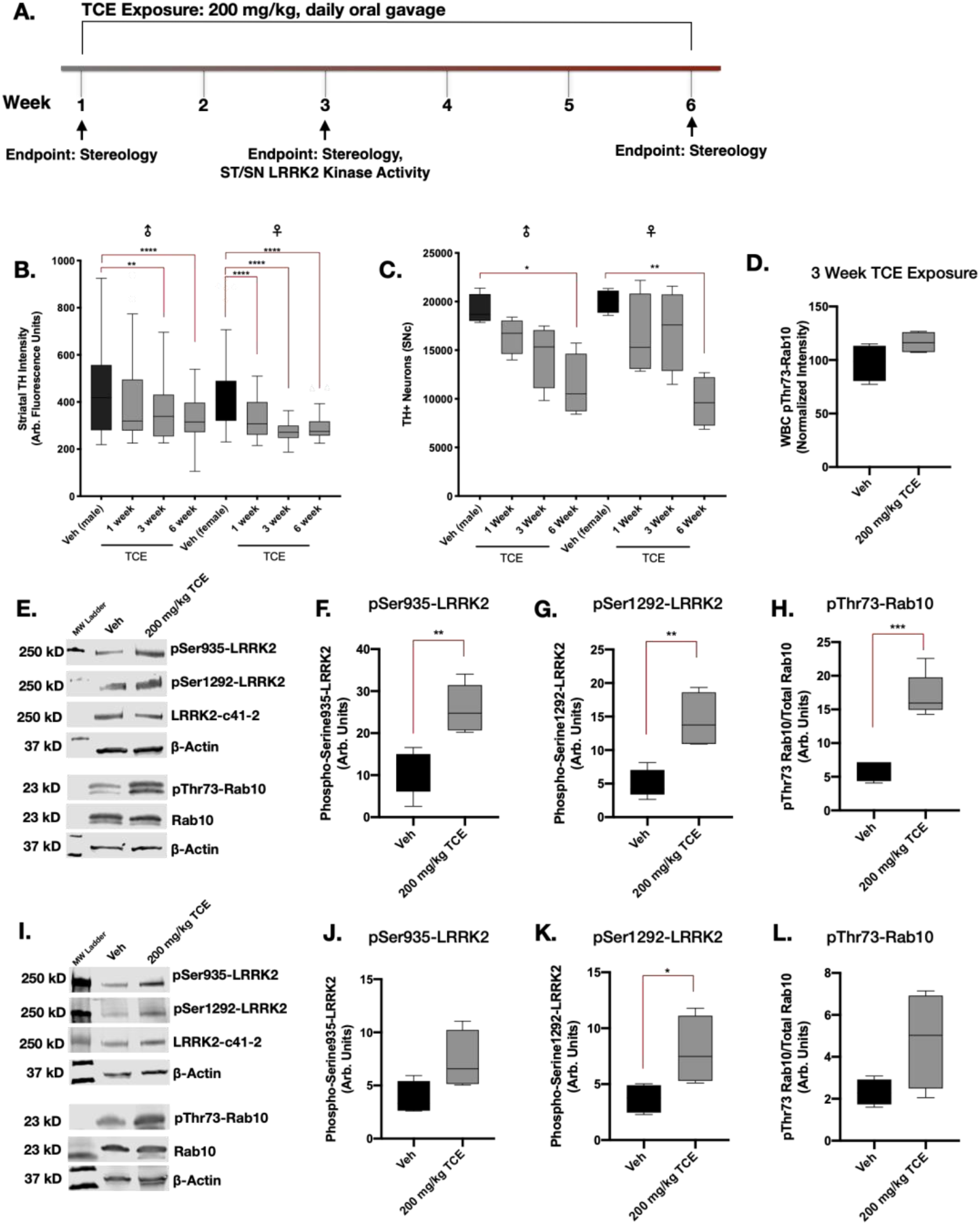
TCE-induced LRRK2 kinase activity is elevated prior to dopaminergic neuron loss. **A.** Aged adult (10 month) male and female Lewis rats were treated with 200 mg/kg TCE or vehicle (olive oil) via daily oral gavage for 1, 3, or 6 weeks. **B-C.** Dopaminergic neuron terminals in the striatum were significantly reduced in male (*p* = 0.0054) and female (*p*<0.0001) animals after 3 weeks of exposure (F(7, 705 = 20.33), *p*<0.0001), however, loss of dopaminergic neurons measured using stereology did not reach significance in male (*p* = 0.005) and female (*p* = 0.0039) rats until 6 weeks of continuous TCE exposure (F(3, 24 = 0.639), *p*<0.0001); two-way ANOVA, N = 5 **D.** pThr73-Rab10 accumulation was not present in cultured WBC of male Lewis rats following 3 weeks of TCE treatment (F(3, 3 = 2.808), *p* = 0.11). **E-H.** Western blot quantification of LRRK2 phosphorylation in the striatum of aged adult (10 month) male Lewis rats after 3 weeks of 200 mg/kg TCE exposure revealed significant elevation in pSer935 (F(4, 4 = 1.168), *p* = 0.0028), pSer1292 (F(3, 4 = 3.924), *p* = 0.003), and pThr73-Rab10 (F(4, 3 = 4.488), *p* = 0.0004). **I-L.** Western blot quantification of LRRK2 phosphorylation in the SN following 3 weeks of TCE exposure shows a significant elevation in pSer1292-LRRK2 (F(3, 3 = 5.820), *p* = 0.0439), but not pSer935 (F(3, 3 = 3.213), *p* = 0.0671) nor pThr73-Rab10 (F(3, 3 = 13.93), *p* = 0.0822); unpaired Τ-test (N = 5), error bars represent SEM.

Quantification of LRRK2 phosphorylation at Ser935, Ser1292, and its kinase substrate Rab10 at pThr73 were measured via western blot in the striatum and SN of male rats after three weeks of daily 200 mg/kg TCE exposure. In the striatum, significantly elevated levels of phosphorylated pSer935-LRRK2 (*p* = 0.0028), pSer1292-LRRK2 (*p* = 0.003), and pThr73-Rab10 (*p* = 0.0004) were observed in TCE treated rats compared to vehicle (Figure 5d-g). Within the SN, three weeks of TCE exposure resulted in elevated pSer1292-LRRK2 (*p* = 0.0439), but not pSer935-LRRK2 (*p* = 0.0671), nor pThr73-Rab10 (*p* = 0.0822; Figure 5h-k). Thus, LRRK2 kinase activity appears to be elevated in nigrostriatal tissue prior to measurable dopaminergic neuron loss. The elevation of pThr73-Rab10 observed in striatal brain tissue after three weeks of TCE treatment did not correlate with elevated pThr73-Rab10 in blood cells at the same timepoint (*p*<0.34; Figure 5d).

### Systemic TCE exposure dose-dependently elevates LRRK2 kinase activity in the rat brain

To further understand the extent of wildtype LRRK2 kinase activity in the brain as a response to systemic TCE exposure, we administered increasing doses of TCE (50, 100, 200 400, or 800 mg/kg) or vehicle (olive oil) for three weeks, via daily oral gavage to separate groups of adult male Lewis rats (Cohort 3). Western blot analysis for pSer935-LRRK2, pSer1292-LRRK2, and pThr73-Rab10 in the striatum and SN of rat brain lysates showed that TCE dose-dependently elevated LRRK2 phosphorylation (Figure 6a-e). As toxicant-induced, wildtype LRRK2 kinase activity in the brain has not been extensively characterized, we compared LRRK2 kinase activity levels from TCE treated rats to whole brain lysates of untreated LRRK2-G2019S knock-in or wildtype mice (Figure 6f-g). As previously reported^39^, LRRK2-G2019S mice displayed significant elevation in pSer1292-LRRK2 (*p* = 0.04) and pThr73-Rab10 (*p* = 0.017), but not pSer935-LRRK2 (*p* = 0.314). Together these data indicate that elevated phosphorylation of wildtype LRRK2, as a measure of increased LRRK2 kinase activity in the brain, is influenced by the relative amount of systemic TCE exposure sustained over a three-week time period.

**Figure 6.**
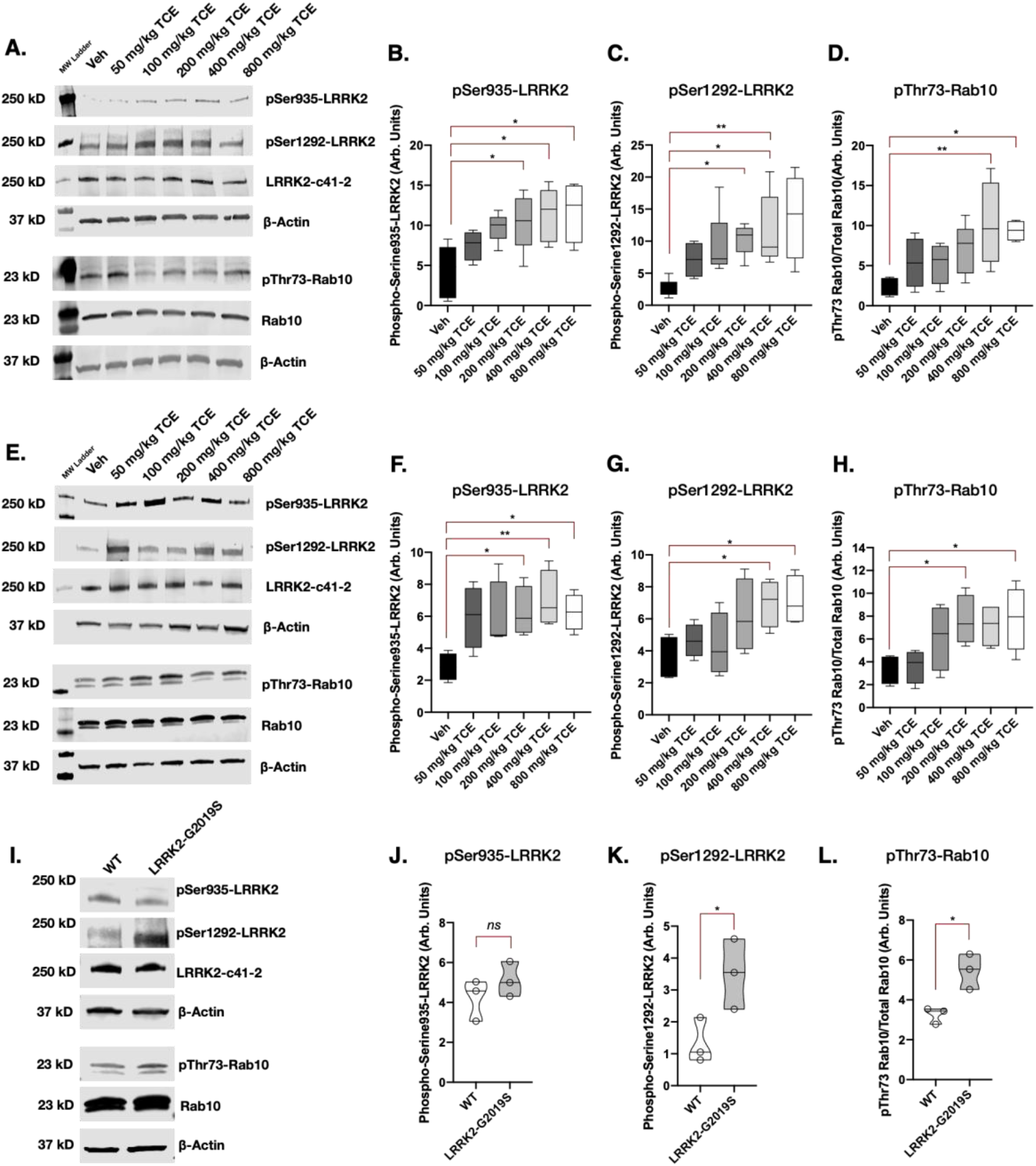
Systemic TCE exposure dose-dependently elevates LRRK2 kinase activity in the rat brain. **A-D.** Western blot analysis of LRRK2 kinase activity in striatal tissue in response to increasing concentrations of TCE (50 – 800 mg/kg) or vehicle (olive oil) given to aged adult (10 month) male Lewis rats for 3 weeks via daily oral gavage; pSer935 (F(5, 18 = 3.806), *p* = 0.0158), pSer1292 (F(5, 22 = 3.3692), *p* = 0.0141), pThr73-Rab10 (F(5, 21 = 4.111), *p* = 0.0093). **E-H.** Corresponding western blot analysis of LRRK2 kinase activity in the SN; pSer935 (F(5, 18 = 3.098), *p* = 0.0344), pSer1292 (F(5, 18 = 3.201), *p* = 0.0306), pThr73-Rab10 (F(5, 18 = 3.385), *p* = 0.0249); one-way ANOVA, N = 5, error bars represent SEM. **I-L.** A comparison of LRRK2 kinase activity in LRRK2-G2019S knock-in mouse brain tissue; pSer935 (F(2, 2 = 1.371), *p* = 0.314), pSer1292 (F(2, 2 = 2.421), *p* = 0.0449), pThr73-Rab10 (F(2, 2 = 4.590), *p* = 0.0176); unpaired Τ-test, N = 3, error bars represent SEM.

## Discussion

There is growing concern for environmental exposures in the etiology for idiopathic PD, as industrialization around the world, particularly in developing countries, is paralleled by the rise of PD incidence^40^. While the link between certain pesticides and PD risk has been extensively characterized^15,41–43^, many industrial byproducts with established neurotoxic properties have not been thoroughly assessed in the context of PD. TCE is a prototypical compound in the class of halogenated solvents, which is comprised of chlorinated, fluorocarbon, brominated or iodinated solvents and their structural derivatives^44^. Thus, while TCE represents one of many industrial solvents, its mechanisms of dopaminergic neurotoxicity may be relative to numerous structurally similar, ubiquitous environmental toxicants.

Previous studies found that high doses of TCE (800 – 1000 mg/kg) in young adult rats or mice produced selective neurodegeneration of dopaminergic neurons, providing proof-of-concept evidence for dopaminergic neurotoxicity from TCE exposure^12,13^. The US Occupational Safety and Health Administration (OSHA) sets daily exposure limits for TCE at 100 ppm over 8 hours, or 300 ppm for five-minute intervals^45^. In contrast, the US Environmental Protection Agency (EPA) sets water limits for TCE at just 5 ppb, however as TCE is classified as a Group 1 carcinogen, there is probably no safe exposure level for TCE in drinking water^44^. At standard temperature and atmospheric pressure, the conversion of ppm to mg/kg for TCE is approximately a 1:1 ratio^46^, therefore, we utilized a dose of 200 mg/kg in this study to model a more moderate level of chronic TCE exposure over 6-weeks.

As aging represents the most significant risk for PD development^47^, we incorporated this factor by using 15-month-old male Lewis rats for the majority of pathological analyses in this study. Indeed, despite receiving a much lower dose of TCE than in previous reports^13^, aged rats used here displayed significant and selective loss of dopaminergic neurons and their striatal terminal projections, corresponding to a reduction in motor behavior (**Figure 1**). Likewise, the death of nigrostriatal dopaminergic neurons appeared to be a progressive, slowly developing retrograde lesion; 10-month-old female and male Lewis rats showed loss of striatal dopaminergic terminals three weeks prior to dopaminergic soma death within the SN during continuous TCE exposure (**Figure 5**). In contrast to other neurotoxicant models^34,48^, no significant sexual dimorphism relating to nigrostriatal neurodegeneration was apparent, perhaps due to the chronic time course over which dopaminergic neurodegeneration occurred. Together, these data demonstrate age, in combination with TCE treatment, is an important co-factor in dopaminergic neurodegeneration, and should be considered in the translation to human risk for PD stemming from past or ongoing solvent exposures.

The predominant risk factor evaluated by this study, however, was the elevation of endogenous, wildtype LRRK2 kinase activity in the brain following systemic exposure to TCE. We demonstrated that TCE treatment induced LRRK2 kinase activation in the nigrostriatal tract, with temporal relevance to a developing dopaminergic lesion (**Figure 5**). In addition, TCE dose-dependently elevated wildtype, endogenous LRRK2 kinase activity, indicating that LRRK2 activation might correlate with the severity of neurotoxicity elicited by exogenous chemicals (**Figure 6**). Other mitochondrial toxicants, such as the pesticide rotenone, similarly activate LRRK2 in dopaminergic neurons (Rocha et al., 2019; Di Maio et al., 2018), indicating that LRRK2 activation is a common theme of PD-associated environmental contaminants. In opposition, reducing LRRK2 kinase activity, either by genetic ablation (LRRK2 knockout) or pharmacological inhibition, seems to prevent or reduce dopaminergic neuropathology caused by environmental toxicant exposure^49–52^. Collectively, these data show that PD risk could be influenced by a gene-environment interaction between LRRK2 and environmental mitochondrial toxicants like TCE.

The mechanism of LRRK2 kinase activation by TCE is less clear. As oxidative stress appears to influence LRRK2 kinase activity^21,53^, the robust oxidative response observed in surviving dopaminergic neurons of TCE treated rats (**Figure 2**) provides a putative source for LRRK2 activation in these cells. For example, Mamais et al.^21^ propose that oxygen radicals indirectly activate LRRK2 by modulating protein phosphatases (e.g. PP1) that dephosphorylate serine residues (Ser910/935) required for 14-3-3 binding, resulting in LRRK2 complex formation at membranes. In this same vein, as LRRK2 membrane association appears to drive its activation status^54,54–56^, ongoing membrane lipid peroxidation caused by TCE, measured here by 4-hydroxynonenal (**Figure 2**), positions membrane-bound LRRK2 in close proximity to oxygen radicals. Membrane dynamics may be of particular importance in TCE exposure and PD risk, as its solvent properties impart physical changes to the lipid bilayer of cellular membranes, thus providing the anesthetic effects of TCE^57^. Whether TCE interaction at the membrane plays a specific role in LRRK2 kinase activation is yet unknown, but could delineate distinct neurotoxic mechanisms of solvents like TCE from other environmental PD toxicants, such as pesticides.

In addition to oxidative stress, there is a high degree of concordance between toxicant models of PD and the loss of endolysosomal function as predegenerative pathology within dopaminergic neurons^33,49^. Most phenotypes of iPD exhibit dysfunction within ALP^58^, as well as protein accumulation, indicating loss of protein degradation pathways are central to dopaminergic neuron injury^59^. Likewise, as the kinase activity of LRRK2 regulates vesicular trafficking and influences the endolysosomal system, mutations (e.g. G2019S) or activation of wildtype LRRK2 leads to disruptions in ALP^60–63^. As such, the impairment of endolysosomal function observed in dopaminergic neurons following six weeks of TCE treatment (**Figure 3**), may be a direct result of LRRK2 kinase activation^64,65^, ongoing oxidative damage to cellular macromolecules^66^, a byproduct of mitochondrial dysfunction^67^, or more likely, a combination thereof. However, as Rocha et al. recently demonstrated, pharmacological kinase inhibition of LRRK2 by the small molecule PF-360 limited endolysosmal dysfunction in dopaminergic neurons induced by rotenone, which was ultimately neuroprotective against cell death^49^. Together, these data indicate a central role for wildtype LRRK2 and endolysosmal dysfunction in toxicant-induced neurodegeneration, and suggest that LRRK2 inhibition might be a protective strategy against exposure to a broad spectrum of environmental toxicants that cause selective dopaminergic neurodegeneration.

As LRRK2 mutations and quantitative trait loci (QTL) are the most common genetic risk factors for PD^68^, environmental toxicant exposures that induce aberrant LRRK2 kinase activity could ‘tip the balance’ toward a PD phenotype in mutation carriers. Toward this end, we observed that TCE induces wildtype LRRK2 autophosphorylation (pSer1292) to a similar extent as brain tissue from G2019S-LRRK2 knock-in mice (**Figure 6**). In contrast, TCE highly elevated pSer935-LRRK2 phosphorylation in rat brain tissue, which was not observed in G2019S-LRRK2 knock-in mice. These data are consistent with the observation that oxidative stress derived from TCE exposure influences upstream signaling kinase (e.g IκB) and phosphatase (e.g. PP1) proteins that regulate wildtype LRRK2 phosphorylation at Ser910/935^21^. In this sense, aberrant activation of wildtype LRRK2 and its downstream consequences may be somewhat distinct from the G2019S-LRRK2 mutation. Whether toxicant-induced LRRK2 activation has an additive or synergistic effect for LRRK2 mutation carriers is unclear. About 50% of individuals with a LRRK2 mutation are clinically diagnosed with PD, suggesting that additional endogenous or exogenous factors, such as TCE, could contribute to overall PD risk in these populations. Studies to determine susceptibility of the dopaminergic system in LRRK2 mutations to environmental toxicant exposure are currently underway in our lab.

## Acknowledgments

This work was supported by research grants from the National Institutes of Health (NS100744, NS095387 to JTG and K99ES029986 to BRD), the American Parkinson Disease Association (JTG), the Parkinson’s Foundation (BRD), the Michael J Fox Foundation, and the friends and family of Sean Logan (JTG).

---

**Supplemental Figure 1.**
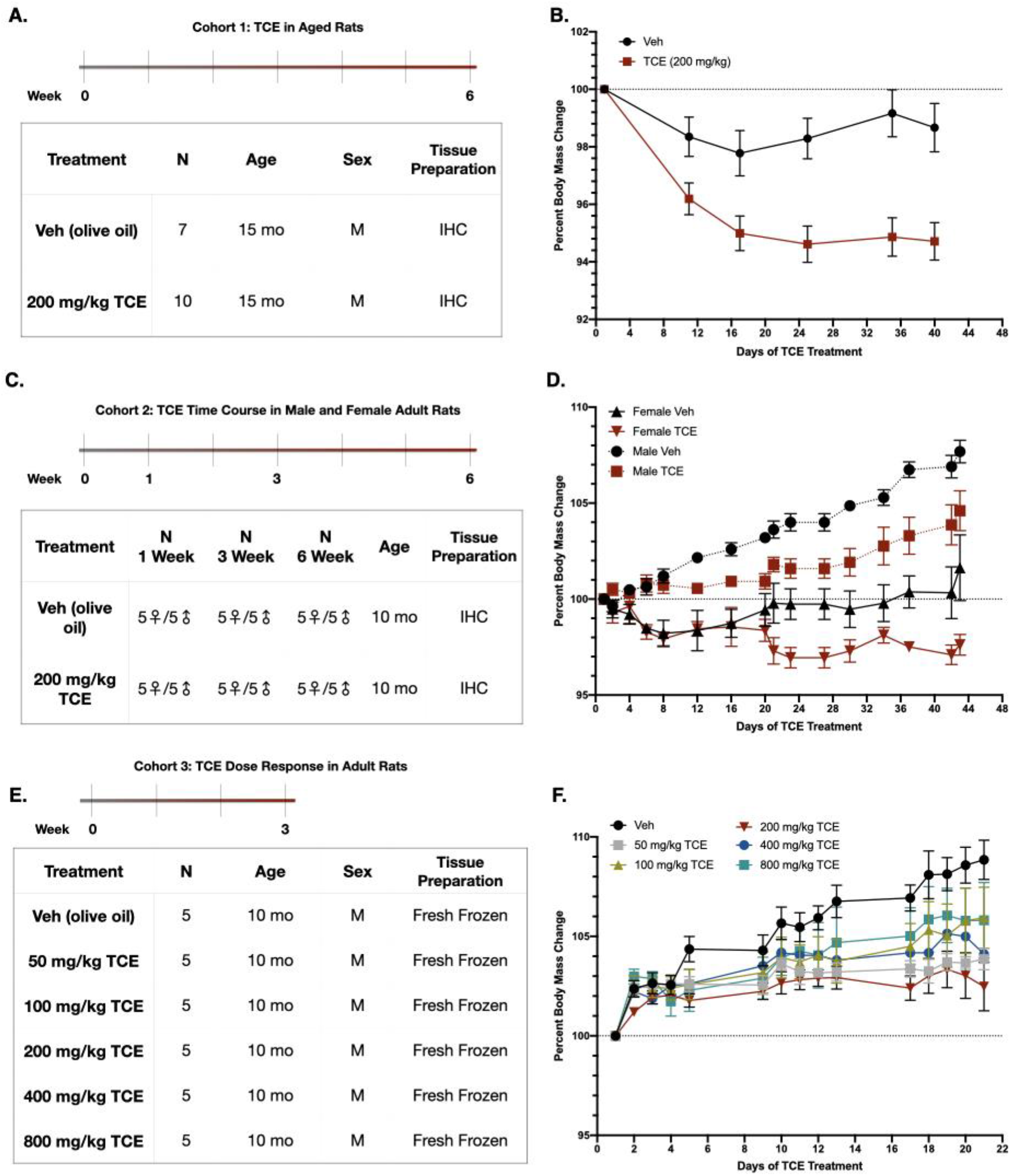
TCE cohort description. **A-B.** 15-Month-old male Lewis rats were treated with 200 mg/kg TCE or vehicle (olive oil) via daily oral gavage for 6 weeks. **C-D.** 10-Month-old male and female Lewis rats were treated with 200 mg/kg TCE or vehicle (olive oil) via daily oral gavage for 1, 3, or 6 weeks. **E-F.** 10-Month-old male Lewis rats were treated with 50, 100, 200, 400, or 800 mg/kg TCE or vehicle (olive oil) for 3 weeks.

**Supplemental Figure 2.**
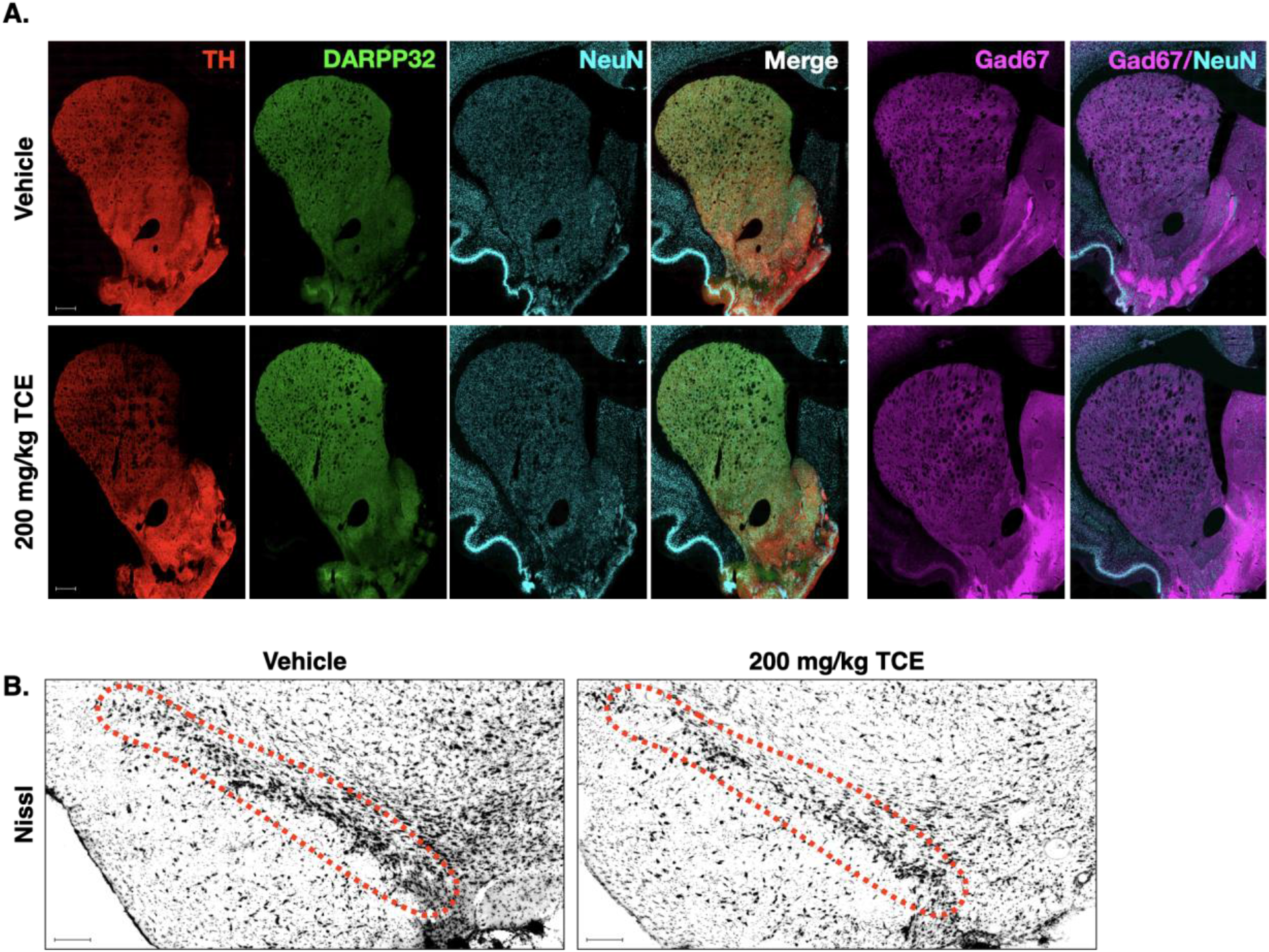
TCE-induced selective degeneration of dopaminergic neurons, terminals. **A.** Representative 20x montage images from the striatum of 200 mg/kg TCE or vehicle treated rats (Cohort 1) demonstrate the loss of TH-positive fibers, but no observable loss in medium spiny neurons (DARPP32, green), total neuron cell bodies (NeuN, cyan), or GABA+ neurons (GAD67, magenta). **B.** Nissl-positive structures within the midbrain of 200 mg/kg TCE or vehicle treated rats demonstrates the loss of cell bodies in conjunction with loss of TH-positive dopaminergic neurons.

